# Social observation influences the trajectory of performance monitoring across trials: evidence from single-trial estimates of the ERN and CRN

**DOI:** 10.1101/2025.09.16.676498

**Authors:** Yanbin Niu, Kianoosh Hosseini, Andy Pena, Carlos Rodriguez, George A. Buzzell

## Abstract

The error-related negativity (ERN) and correct-related negativity (CRN) are event-related potentials (ERPs) that reflect performance monitoring following error and correct responses, respectively. Prior work demonstrates the ERN is sensitive to the motivational significance of errors, which increases under social observation. However, most studies testing how social observation impacts performance monitoring rely on trial-averaged ERPs, potentially obscuring meaningful fluctuations in ERN/CRN over time. Here, we had participants complete a Flanker task twice (social observation vs. alone) and employed mixed-effects modeling of single-trial ERPs to test if social observation impacts ERN/CRN trajectories over short (within blocks) or long (between blocks) timescales. We found that social observation selectively influenced ERN/CRN trajectories over short timescales: for blocks performed under social observation (but not alone), ERN magnitudes increased across trials and CRN magnitudes decreased. At longer timescales, ERN/CRN significantly decreased across all blocks, regardless of social observation and consistent with a vigilance decrement. To our knowledge, this is the first demonstration that social observation influences performance monitoring trajectories over short timescales. Results highlight the importance of analyzing ERN/CRN trajectories over relatively short timescales to fully characterize the impact of social observation on performance monitoring dynamics. These findings lay the groundwork for future investigation into whether social observation interacts with individual differences in motivation/affect to differentially impact performance monitoring dynamics.

## Introduction

Performance monitoring refers to a set of neurocognitive processes that allow individuals to monitor and detect the need to adapt their behavior (Ullsperger, Fischer, Nigbur, & Endrass, 2014).), Performance monitoring is often studied using the high-temporal resolution afforded by event-related potentials (ERPs; Ullsperger et al., 2014). For example, the error-related negativity (ERN) is a negative ERP deflection that emerges over frontocentral scalp regions within 100 ms of making an error—thought to index error detection (i.e., error monitoring; Falkenstein, Hohnsbein, Hoormann, & Blanke, 1991; Gehring, Goss, Coles, Meyer, & Donchin, 1993). The correct-related negativity (CRN) refers to a similar—albeit smaller—negative frontocentral deflection occurring within 100 ms of correct responses (Ford, 1999; F Vidal, Hasbroucq, Grapperon, & Bonnet, 2000). Prior work demonstrates that the ERN is sensitive to the motivational significance of errors (Hajcak, Moser, Yeung, & Simons, 2005a). For example, preforming tasks under social observation, increases ERN magnitude and/or the relative difference between ERN/CRN (e.g., Barker, Troller-Renfree, Bowman, Pine, & Fox, 2018; Buzzell et al., 2017; Clayson et al., 2024; Kim, Iwaki, Uno, & Fujita, 2005; Niu, Li, Pettit, Buzzell, & Zhao, 2023; Voegler et al., 2018). However, prior work investigating impacts of social observation on performance monitoring have almost exclusively relied on trial-averaged ERPs, which may obscure important trial-by-trial dynamics. In the current study, we leveraged single-trial analyses and mixed-effects modeling (Heise, Mon, & Bowman, 2022a; Volpert-Esmond et al., 2018) to investigate whether social observation impacts the trajectory of performance monitoring ERPs at short (trial-level) and long (block-level) timescales. Closing this gap in the literature could inform how social observation impacts performance monitoring within dynamic, real-world social environments.

Prior work demonstrating that social observation enhances the ERN largely mirrors the effects of other motivational factors (e.g., Hajcak et al., 2005), consistent with the notion that social evaluation increases motivation to perform well in healthy adults (Olvet & Hajcak, 2008). However, in at least some studies, effects of social observation manifest as an increase in the relative difference between the ERN/CRN (Clayson et al., 2024; Schillinger, De Smedt, & Grabner, 2016) as opposed to an ERN-specific effect per se. To understand relations between social observation and the ERN/CRN, it is important to distinguish between what each of these ERP components index. Functional interpretations of the ERN and CRN remain under investigation (for reviews, see Gehring, Liu, Orr, & Carp, 2012; Franck Vidal, Burle, & Hasbroucq, 2021). Nonetheless, a larger (more negative) ERN magnitude is generally thought to at least partially involve processing specific to error detection, which scales with the motivational significance of errors. In contrast, a larger (more negative) CRN magnitude may reflect more general aspects of performance monitoring present on correct trials, including increases in uncertainty and/or misclassifying correct trials as errors (Pailing & Segalowitz, 2004; Scheffers & Coles, 2000). Therefore, to the extent that motivational factors drive adaptive increases in task-related attention and improve the fidelity of performance monitoring, error and correct responses should be more accurately signaled by the brain, reflected in a larger (or relatively larger) ERN for errors and a relatively smaller CRN for corrects. Indeed, such a pattern is typically observed when adults and youth perform tasks under social evaluation, with studies reporting increases in the ERN (Hajcak et al., 2005a; Kim et al., 2005; Niu et al., 2023; Voegler et al., 2018) or increases in the relative difference between the ERN/CRN (Clayson et al., 2024; Schillinger et al., 2016). Nonetheless, some discrepancies exist in the literature, which we hypothesize may at least partially be explained by condition-averaged ERPs obscuring nuanced trajectories of ERN/CRN under social observation.

Prior studies investigating the impact of social observation on the ERN have focused on trial-averaged condition differences as opposed to examining trial- or block-level trajectories. Yet, increasing evidence demonstrates the ERN (and CRN) often fluctuate meaningfully over the course of a task. For example, over relatively longer timescales (across blocks), both the ERN and CRN decrease in magnitude as the result of fatigue (vigilance decrements; Boksem et al., 2006; Bonnefond et al., 2011; Clayson, Baldwin, & Larson, 2025; Volpert-Esmond et al., 2018). Here, it is important to emphasize that simultaneous decreases in *both* the ERN and CRN are indicative of fatigue effects, consistent with broad disengagement of task-related performance monitoring. Thus, the functional interpretation of an observed CRN decrease is dependent on concomitant measurement of the ERN. CRN decreases that co-occur with ERN decreases indicate broad disengagement of task-related performance monitoring (e.g., due to fatigue). Conversely, CRN decreases that co-occur with a stable/increasing ERN (i.e., an increase in the relative ERN/CRN difference) indicate enhanced performance monitoring.

Although prior work demonstrates that at least some forms of motivational enhancement (e.g., monetary incentives) partially counteract the effects of fatigue on the ERN (Boksem et al., 2006), no studies have investigated whether social observation impacts the ERN over longer timescales. Preliminary evidence from one relatively small study (N=18 for a between-subjects manipulation) suggests social observation can modulate long-term trajectories of the CRN (Bonnefond et al., 2011). However, as the functional interpretation of the CRN relies on concurrent ERN measurement, work is needed that simultaneously assesses the effects of social observation on trajectories of both ERN and CRN.

To our knowledge, no prior studies have investigated whether social observation impacts trajectories of ERN/CRN over relatively short timescales (i.e., across trials within a block). This is a critical gap, given that effects of social observation on performance monitoring ERPs (ERN/CRN) are typically studied using task paradigms organized into blocks of trials. Blocking trials effectively creates a set of event boundaries that segregate the task into individual, self-contained task units (Zacks et al., 2001; Zacks & Swallow, 2007). Moreover, the break point between blocks can often serve as an opportunity for explicit reminders of social observation, including protocols that involve block-level feedback being provided by the observer (e.g., Barker et al., 2018; Buzzell et al., 2017; Niu et al., 2023). As a result, one possibility is that effects of social observation on ERN/CRN “ramp up” over the course of a block. reflected in a trajectory of increasing (more negative) ERN magnitude and/or relative increases in the ERN/CRN difference leading up to the end of each block. Indirect support for this idea comes from studies demonstrating that the anticipation of social evaluation increases activation within the performance monitoring network (Knutson & Cooper, 2005). An alternative hypothesis is that effects of social observation on performance monitoring would instead be more pronounced earlier within a given block, and then subsequently decrease over time. Regardless, in the absence of data investigating the impact of social observation on trial-level ERN/CRN dynamics, it remains completely unknown as to whether social observation impacts trajectories of performance monitoring at the within-block level.

In the current study, we employed mixed-effects modeling of single-trial ERPs to investigate whether social observation influences the trajectory of ERN/CRN at either short (within blocks) or long (between blocks) timescales. Data were acquired from participants that completed a Flanker task twice: once while under social observation and once while alone. Our hypotheses were as follows. First, we hypothesized that social observation (compared to the nonsocial condition) would elicit an overall increase in ERN magnitude (and/or the ERN/CRN difference). Second, counteracting any potential vigilance decrements present for ERN/CRN over longer timescales (across blocks), we hypothesized that social observation (compared to the nonsocial condition) would serve to either maintain or increase ERN magnitudes (and/or relative ERN/CRN differences) across blocks. Third, we hypothesized that over shorter timescales (across trials within a block), social observation would again serve to either maintain or increase ERN magnitudes (and/or relative ERN/CRN differences) across trials within a given block.

## Methods

### Participants

Sixty healthy adults participated in the study after providing informed consent. Participants received either monetary compensation or course credit for participation. All participants were fluent in English and reported no prior head injury resulting in loss of consciousness. Of the initial 60 participants, one was excluded due to potential familiarity with the confederate and five were excluded due to technical issues. This left 54 participants for behavioral analyses (Mean age = 22.81 years, SD = 3.40; 48 f, 6 m), with 45 of these individuals also having EEG recorded (Mean age = 23.01 years, SD = 3.64; 40 f, 5 m). For EEG analyses, five participants were removed from the nonsocial condition and six from the social condition, due to having less than six artifact-free error trials following preprocessing (Pontifex et al., 2010; Steele et al., 2016). Thus, final EEG data analyses included 40 participants for the nonsocial condition and 39 participants for the social condition.

### Procedures

Participants first completed a battery of surveys to assess demographic information and individual differences in thoughts and feelings. Following a practice session, participants completed a Flanker task twice (in a counterbalanced order) while EEG was recorded. In the social observation condition, participants were observed by a second experimenter via Zoom and using a stand-mounted iPad positioned just behind and to the right of the participant (out of direct view). The participant was informed that the second experimenter would observe their performance throughout the social condition, as well as read the block feedback displayed on the participant’s computer screen after each block. In the nonsocial condition, the iPad was turned off and covered with a sheet and participants were explicitly informed that they would complete the task alone and without observation (see Figure 1A). The same block feedback was presented on the participant’s screen after each block in the nonsocial condition. Following completion of the Flanker tasks, participants engaged in additional computer and social-interaction tasks; results of these additional tasks are beyond the scope of the current report and not discussed further.

**Figure 1.**
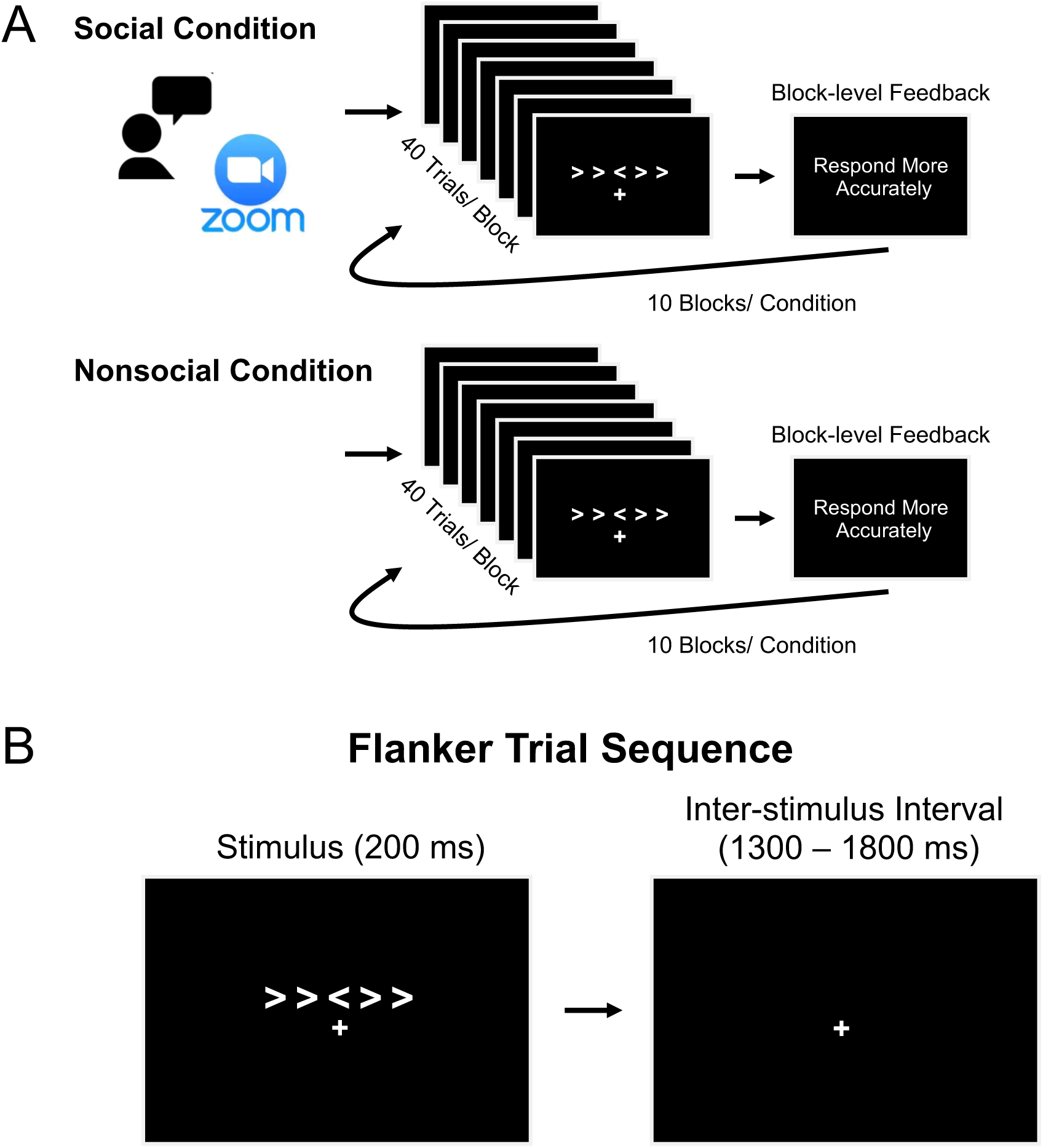
Task structure and trial sequence. A. In the social condition, participants completed the Flanker task while being observed via Zoom by an experimenter who provided block-level feedback. In the nonsocial condition, the same task and block-level feedback were presented, but participants were explicitly told they were completing the task alone. Each condition consisted of 10 blocks of 40 trials, with feedback presented at the end of each block based on task performance. B. Flanker trial sequence. Each trial began with a fixation cross and presentation of a five-arrow array for 200 ms. Participants responded to the direction of the central arrow. The inter-stimulus interval varied randomly between 1300 and 1800 ms.

### Flanker task

Participants completed a modified Flanker task (Eriksen & Eriksen, 1974). On each trial, an array of five arrows was presented and participants were required to respond to the direction of the central arrow while ignoring the flanking arrows. Participants used their right/left thumbs to indicate the right/left direction of the target arrow via button box (The Black Box ToolKit Ltd., Sheffield, UK). Flanking arrows could either be congruent (pointing in the same direction as the target) or incongruent (pointing in the opposite direction of the target). During each trial, the arrows appeared at trial start and remained onscreen for 200 ms, followed by an inter-stimulus interval randomly jittered from 1300-1800 ms (see Figure 1B). Participant responses were recorded throughout the duration of each trial. Throughout each block, a fixation cross remained at the center of the screen, just below the location where the array of arrows appeared on each trial. Within each condition (social, nonsocial) participants completed 10 blocks of 40 trials (400 trials/condition), with an equal mix of congruent/incongruent trials per block. To maintain appropriate error rates, block-level feedback was presented onscreen after each block (Gehring et al., 2012). If accuracy was between 75% and 90%, “Good job” was displayed. When accuracy exceeded 90% or fell below 75%, “Respond faster” or “Respond more accurately” were displayed, respectively. The experiment was conducted on a 15-inch Lenovo Legion 7i laptop running Windows 10, using PsychoPy version 2021.2.3 (Peirce et al., 2019) to present the stimuli.

### EEG acquisition and preprocessing

64-channel EEG was collected at 1000 Hz via a 64-channel EasyCap custom EEG cap (EasyCap GmbH, Herrsching, Germany; see Figure 2 for cap layout) and a Brain Products actiCHamp amplifier using BrainVision Recorder software (Brain Products GmbH, Munich, Germany). Impedance was reduced to a targeted level of ≤ 25 kΩ prior to data collection. EEG electrodes were referenced to electrode 1 (∼FCz) during recording and re-referenced to the average during preprocessing. EEG data were preprocessed using MATLAB R2021b (MathWorks Inc., Sherborn, MA, USA), the EEGLAB toolbox, and a modified version of the MADE pipeline (Debnath et al., 2020; Delorme & Makeig, 2004). Complete preprocessing details are reported in the Supplementary Materials. Data were segmented into 3-second epochs (-1 to 2 seconds, relative to the response) and the (-400 to -200) pre-response baseline was subtracted from each epoch.

**Figure 2.**
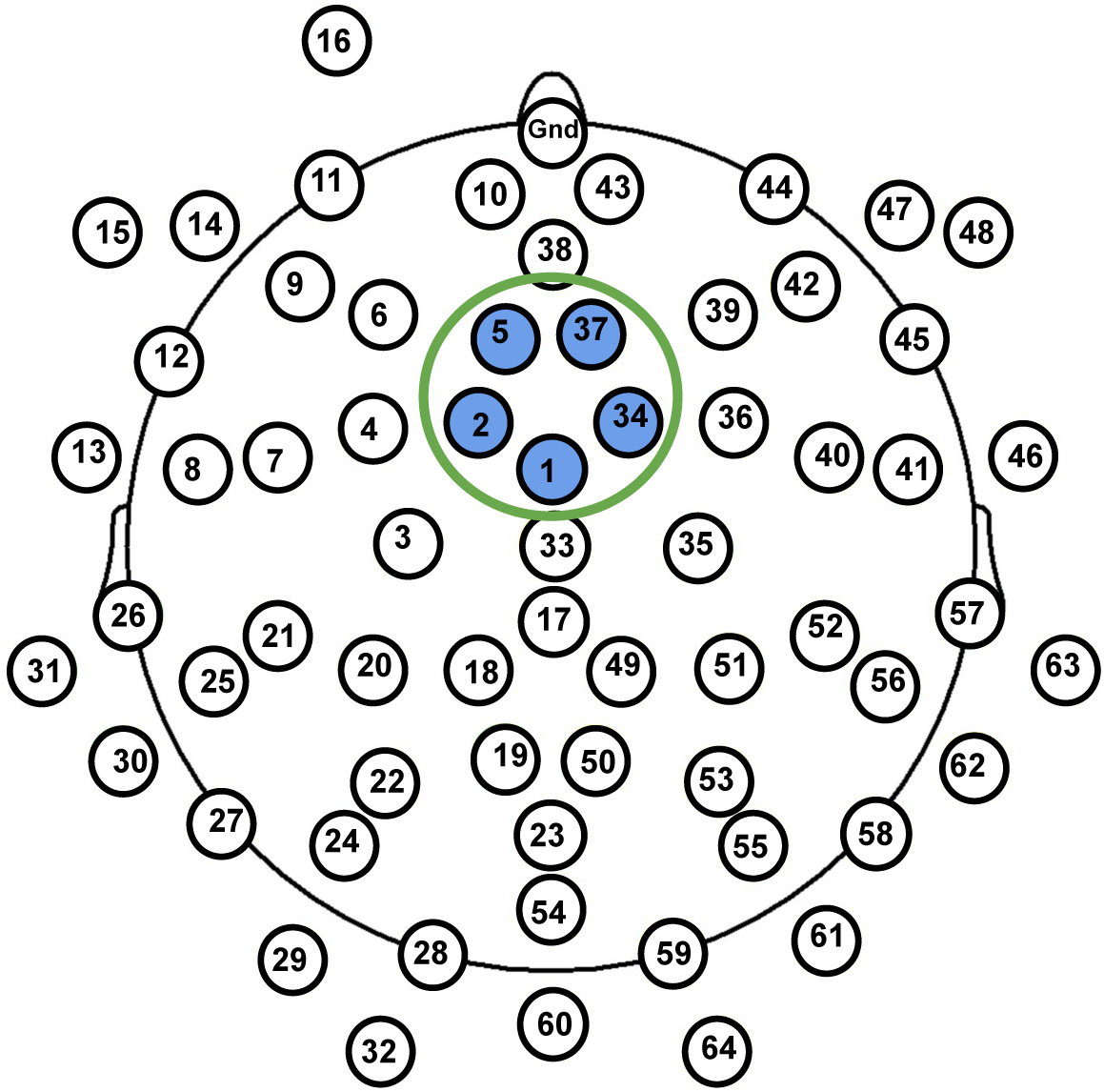
64-channel EasyCap EEG cap layout. Note. The selected electrodes over the frontocentral electrode cluster (E1, E2, E5, E34, E37) are shown in the blue circle. Note that this cluster approximates the location of FCz and surrounding electrodes in a standard 10-10 system.

### Flanker task behavior

To be included in the behavioral and EEG analyses, participants were required to have an overall accuracy rate greater than 70%; all participants met this criterion (mean accuracy rate = 87.39% for the nonsocial condition, mean accuracy rate = 86.95% for the social condition). Due to the positive skew typically observed in RT data, all RT values were log-transformed following standard procedures (Luce, 1986) prior to performing statistical analyses; results are also reported in the aggregate as raw RT for ease of interpretation.

### ERN/CRN

Only incongruent trials were included in analyses of ERN/CRN to isolate error-related effects and minimize stimulus-related confounds, as errors are more prevalent in incongruent trials in the Flanker task (Buzzell et al., 2019; Eriksen & Eriksen, 1974). Thus, incongruent-error trials were used to measure the ERN, and incongruent-correct trials used to measure the CRN. For the remainder of the manuscript, we refer to incongruent-error and incongruent-correct trials as “error” and “correct” for simplicity. At a frontocentral electrode cluster (E1, E2, E5, E34, E37; see Figure 2), where error-related brain activity is maximally negative (Gehring et al., 2012; Meyer, Mehra, & Hajcak, 2021), trial-level mean amplitudes for the CRN and ERN were extracted within the 0–100 ms window following correct and incorrect responses, respectively.

### Analytic plan

All statistical analyses were conducted using R v4.3.3 (R Core Team, 2013). Prior to analysis, continuous variables and discrete numerical variables (block and trial number) were standardized (z-scored), and binary predictor variables (accuracy and observation condition) were factorized and contrast coded (-1 and 1). Linear Mixed Models (LMMs) with a continuous outcome variable were fit using the ‘lme’ function from the ‘nlme’ package (version 3.1-166; Pinheiro, Bates, & R Core Team, 2022). Generalized Linear Mixed Models (GLMMs) with a binary outcome variable were fit using the ‘glmer’ function from the ‘lme4’ package (version 1.1-35.5; Bates, Mächler, Bolker, & Walker, 2015). In all models, subject was entered as random intercept. Where appropriate, significant interaction effects were probed via the ‘emmeans’ and ‘emtrends’ functions from the ‘emmeans’ package (version 1.10.2; Lenth, 2022).

### Preliminary analyses

We began with preliminary analyses of subject-averaged behavioral data to confirm expected effects of congruency on accuracy rates and RT. For these analyses, separate LMMs were conducted for accuracy rates and RT. The model predicting log-transformed RTs included accuracy, congruency, observation condition; the model predicting accuracy rates included congruency and observation condition, and their interactions.

### Hypothesis testing

We had three hypotheses: 1) social observation would elicit an overall larger ERN (or ERN/CRN difference) compared to a nonsocial condition; 2) at longer timescales (across blocks), social observation would serve to either maintain or increase ERN magnitude (or ERN/CRN differences) compared to a nonsocial condition; 3) at shorter timescales (across trials within a block), social observation would either maintain or increase ERN magnitude (or relative ERN/CRN differences) compared to a nonsocial condition. For parsimony, we jointly tested our three hypotheses by fitting a single LMM predicting trial-level ERP amplitude (ERN/CRN) from accuracy, observation condition, block number, and trial number (within a block), along with all relevant interactions.

### Exploratory analyses

To examine whether any identified patterns observed in the trial-level ERN/CRN data were also present in behavior, we conducted a parallel set of analyses predicting trial-level behavior. These exploratory analyses were restricted to incongruent trials, matching the ERP analysis approach. We fit an LMM predicting trial-level log-transformed RTs from accuracy, observation condition, block number, and trial number (within a block), along with all relevant interactions. Similarly, we fit a GLMM predicting trial-level accuracy-outcomes from observation condition, block number, and trial number (within a block), along with relevant interactions.

We additionally fit two exploratory LMMs to test whether block-level feedback or trial-level RT could account for any observed effects reported in our primary, trial-level ERN/CRN analyses. First, we tested whether the type of block-level feedback presented before each block modulated the trial-level ERP effects reported in our primary analyses. This model included accuracy, observation condition, trial number (within a block), and feedback type, along with their interactions. Second, we examined whether trial-level behavioral responses (i.e., RT) were modulated the trial-level ERP effects reported in our primary analyses by fitting a similar model and swapping out the feedback type predictor for log-transformed RTs.

## Results

### Subject-level behavioral data

Consistent with prior Flanker task studies (Eriksen & Eriksen, 1974), subject-level analyses of accuracy rates revealed a significant main effect of congruency (β = -0.701, 95% CI [-0.770, -0.631], *p* < 0.001), with participants responding more accurately on congruent trials (Mean = 94.87%, SE = 0.92%) compared to incongruent trials (Mean = 79.47%, SE = 0.92%). There was neither a significant main effect of social condition (*p* = 0.573), nor a significant congruency-by-condition interaction (*p* = 0.239).

Analyses of subject-level RT data also revealed expected effects, including a significant main effect of congruency (β = 0.236, 95% CI [0.176, 0.295], *p* < 0.001), with participants responding faster on congruent trials (Mean log RT = 5.92 (372ms), SE = 0.025) compared to incongruent trials (Mean log RT = 6.03 (416ms), SE = 0.024). Additionally, the expected main effect of accuracy on RT was significant (β = -0.340, 95% CI [-0.399, -0.281], *p* < 0.001), with participants responding faster on error trials (Mean log RT = 5.89 (361ms), SE = 0.025) compared to correct trials (Mean log RT = 6.06 (428ms), SE = 0.024). A significant congruency-by-accuracy interaction was also observed (β = -0.124, 95% CI [-0.183, -0.065], *p* = 0.001). On average, participants responded faster on correct congruent trials (Mean log RT = 5.97 (392ms), SE = 0.027) compared to correct incongruent trials (Mean log RT = 6.14 (464ms), SE = 0.027), whereas there was no difference in RT between error congruent trials (Mean log RT = 5.86 (351ms), SE = 0.027) and error incongruent trials (Mean log RT = 5.92 (372ms), SE = 0.027). There was no significant main effect of social condition (*p* = 0.066), nor a significant congruency-by-accuracy-by-condition interaction (*p* = 0.286).

### Trial-level ERN/CRN as a function of observation condition, trial, and block

Consistent with with prior work (Niu et al., 2023; Voegler et al., 2018), we observed the expected main effect of accuracy (β = 0.257, 95% CI [0.235, 0.279], *p* < 0.001), with ERP responses being more negative on error (Mean = -2.760, SE = 0.352) compared to correct (Mean = 0.559, SE = 0.331) trials; see Figure 3. Additionally, there was a significant positive association between block number and ERP responses (β = 0.024, 95% CI [0.002, 0.045], *p* = 0.029), indicating that as the blocks progressed, ERN/CRN amplitudes became smaller (i.e., less negative), consistent with a vigilance decrement observed in prior work (Boksem et al., 2006; Volpert-Esmond et al., 2018). Contrary to our hypotheses, we did not observe an overall effect of social observation, as the main effect of observation condition was not significant (*p* = 0.556). Similarly, there was no evidence that social observation influenced trajectories of ERN/CRN at longer timescales (across blocks), as all interactions between observation condition and block number were not significant (all *p* > .470). Nonetheless, we observed a significant three-way interaction between accuracy, observation condition, and trial number within a block (β = 0.033, 95% CI [0.011, 0.054], *p* = 0.003), consistent with our hypothesis that social observation impacts trajectories of ERN/CRN over shorter timescales (across trials within a block); see Figure 4. The nature of this interaction was such that, in the social condition, the ERN significantly increased in magnitude (became more negative) as the number of trials progressed within a block (β = -0.061, 95% CI [-0.114, -0.007], *p* = 0.026), whereas the CRN became significantly smaller in magnitude (more positive) across trials within a block (β = 0.041, 95% CI [0.018, 0.064], *p* = 0.001). Moreover, simple slopes for ERN and CRN across trials (within a block) significantly differed from each other within the social condition (t = -0.102, SE = 0.030, *p* = .004). In contrast, in the nonsocial condition, no significant changes across trials (within a block) were detected for either the ERN (*p* = 0.399) or CRN (*p* = 0.731), and their simple slopes also did not significantly differ (*p* = .799; see Figure 4). Full model results are presented in Table 1.

**Figure 3.**
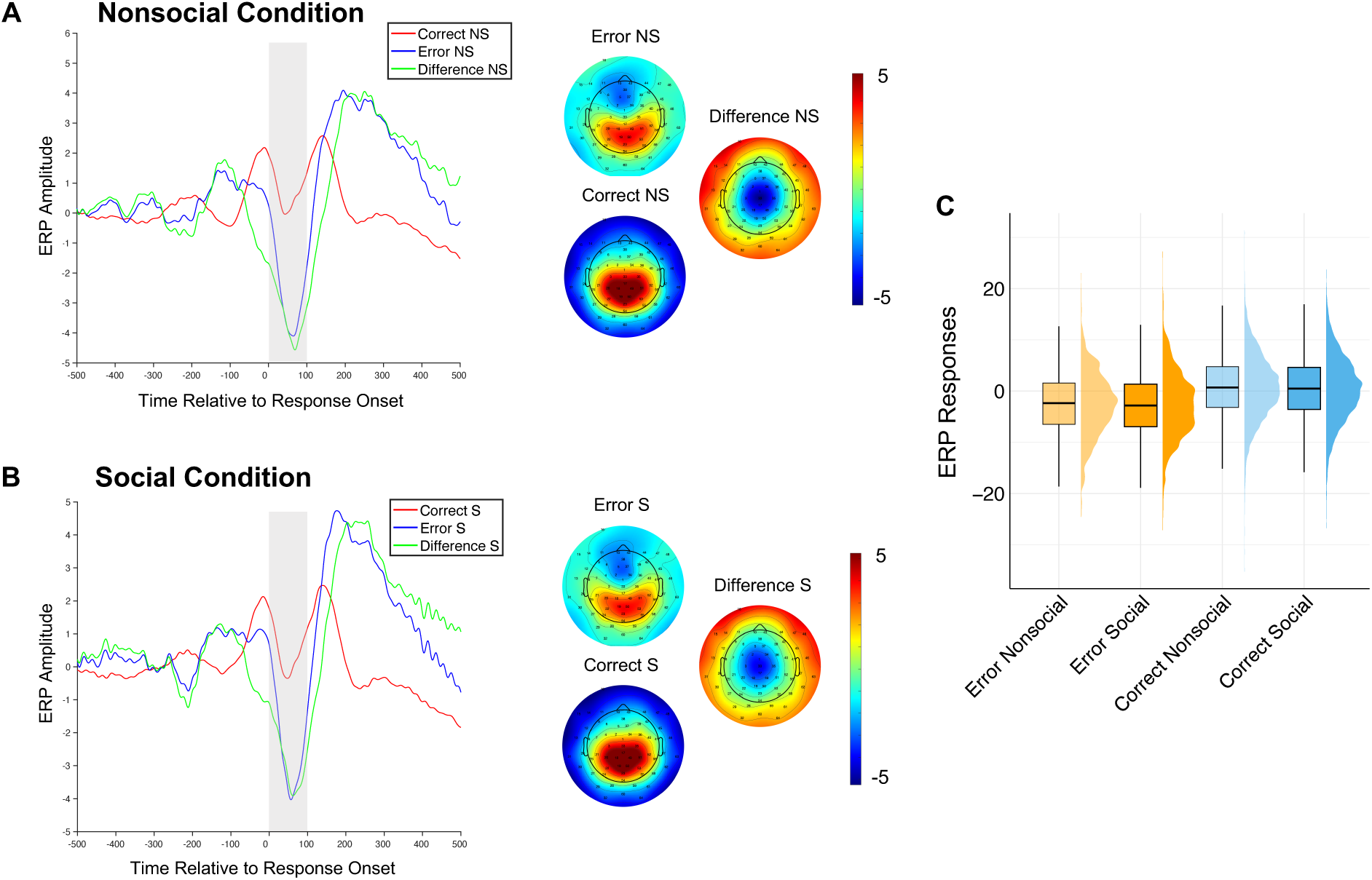
ERP responses by accuracy and observation condition. A. Grand average ERP waveforms at the frontocentral cluster (FCz and surrounding electrodes) in the nonsocial condition (NS) for correct (blue) and error (red) trials, along with the error–correct difference wave (green). Corresponding scalp topographies are shown for each trial type and their difference. B. ERP waveforms and topographies for the social condition (S), plotted using the same conventions. Shaded regions in the time-course plots represent the window used to extract mean ERP amplitudes. C. Violin plots showing the distribution of trial-level ERP amplitudes by trial type (correct vs. error) and observation condition (nonsocial vs. social).

**Figure 4.**
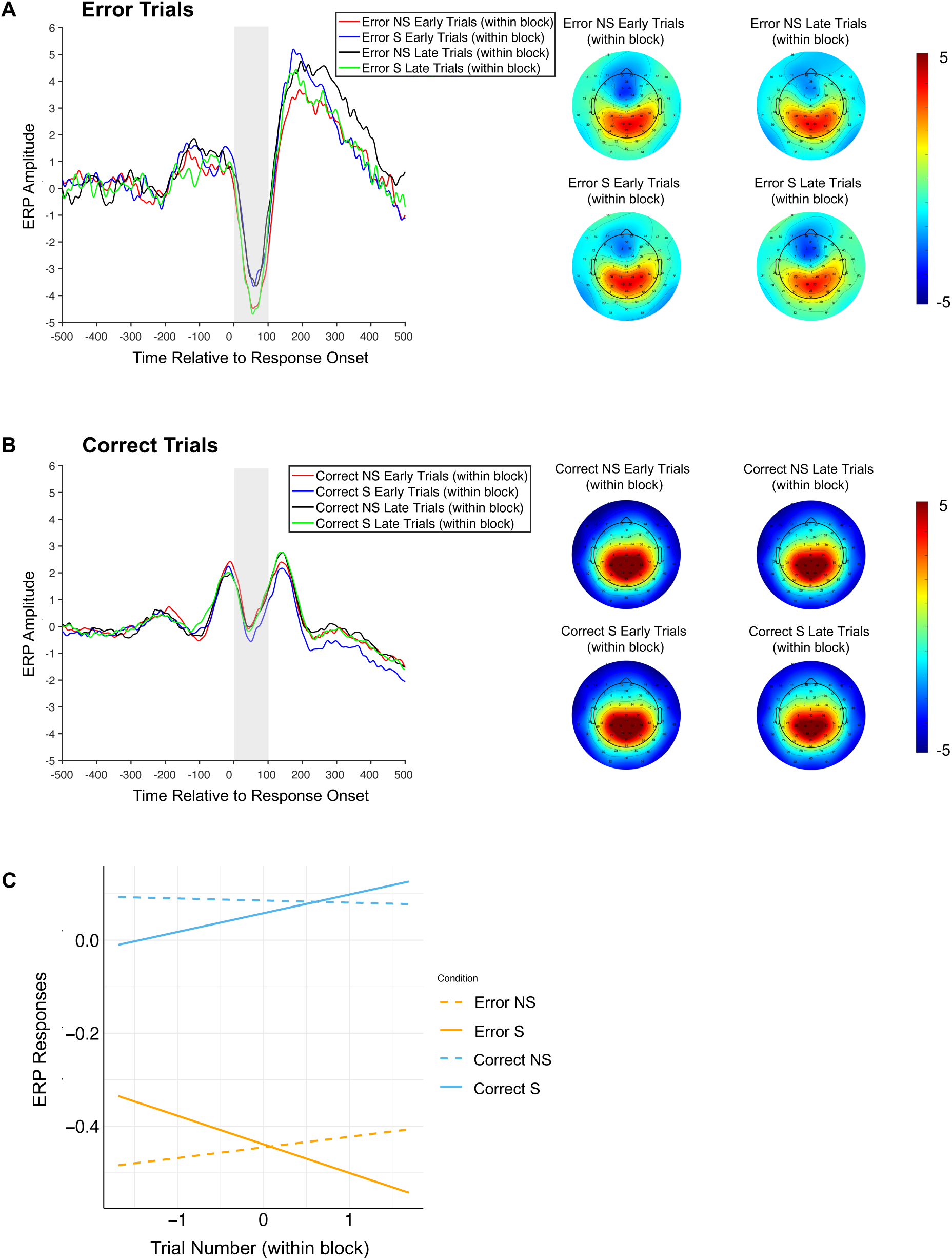
ERP responses across early and late trials within blocks. A. Grand average ERP waveforms for error trials, separated by observation condition: nonsocial (NS), social (S). Statistical analysis incorporated all trials; for visualization purposes only, early trials were defined as trials 1–13 and late trials as trials 28–40 within each 40-trial block, averaged across blocks. Corresponding scalp topographies are shown for each condition and trial grouping (display purposes only). B. ERP waveforms and topographies for correct trials, plotted using the same conventions. Shaded areas in the waveforms indicate the time window used to extract mean ERP amplitudes. C. Model-predicted ERP amplitudes plotted as a function of trial number within a block, separated by response type and observation condition.

**Table 1.**
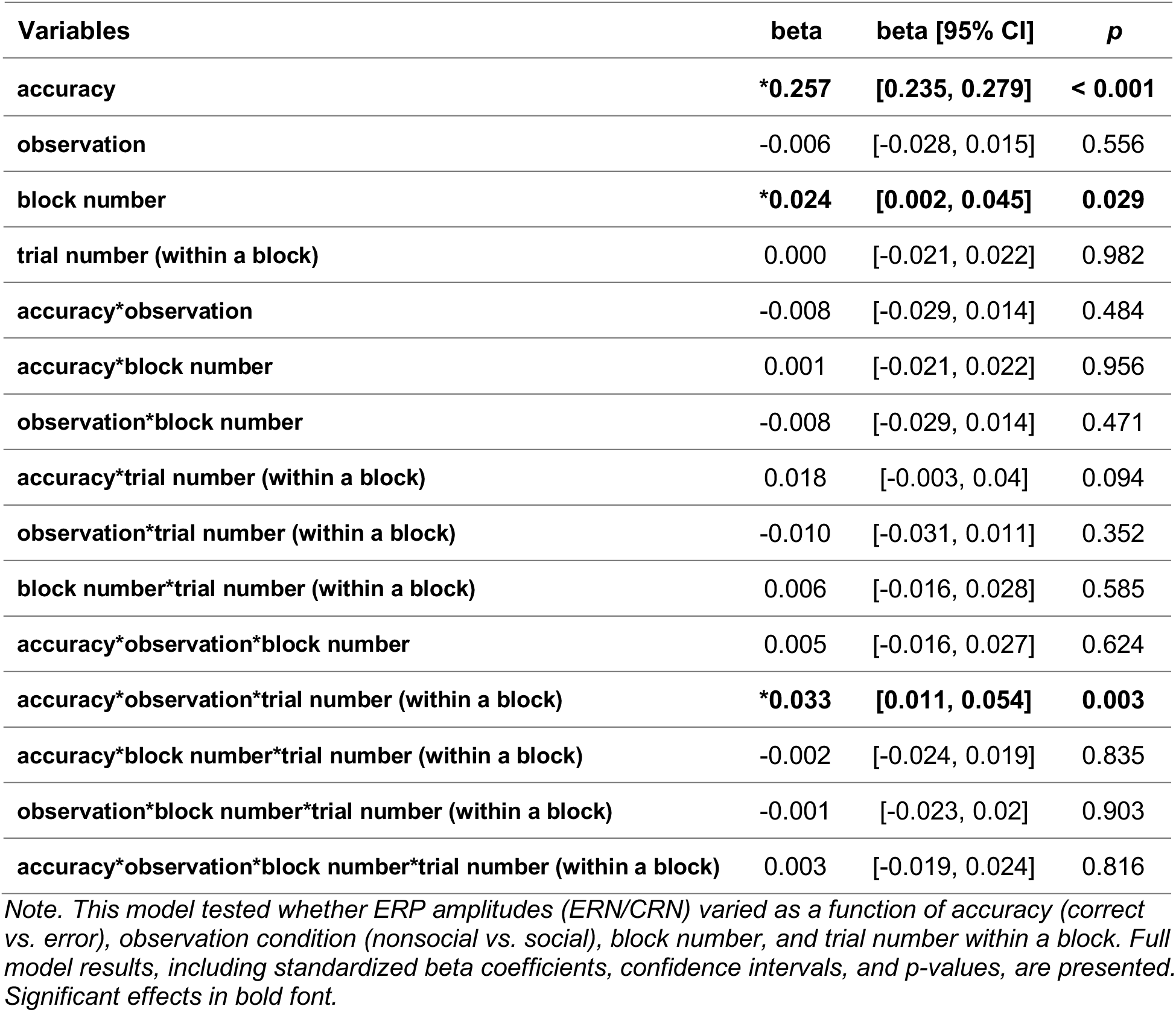
Linear mixed-effects model predicting trial-level ERN/CRN from accuracy, observation condition, block number, and trial number within a block.

|  |  |  |  |
| --- | --- | --- | --- |
| **accuracy** | **\*0.257** | **[0.235, 0.279]** | **< 0.001** |
| **observation** | -0.006 | [-0.028, 0.015] | 0.556 |
| **block number** | **\*0.024** | **[0.002, 0.045]** | **0.029** |
| **trial number (within a block)** | 0.000 | [-0.021, 0.022] | 0.982 |
| **accuracy\*observation** | -0.008 | [-0.029, 0.014] | 0.484 |
| **accuracy\*block number** | 0.001 | [-0.021, 0.022] | 0.956 |
| **observation\*block number** | -0.008 | [-0.029, 0.014] | 0.471 |
| **accuracy\*trial number (within a block)** | 0.018 | [-0.003, 0.04] | 0.094 |
| **observation\*trial number (within a block)** | -0.010 | [-0.031, 0.011] | 0.352 |
| **block number\*trial number (within a block)** | 0.006 | [-0.016, 0.028] | 0.585 |
| **accuracy\*observation\*block number** | 0.005 | [-0.016, 0.027] | 0.624 |
| **accuracy\*observation\*trial number (within a block)** | **\*0.033** | **[0.011, 0.054]** | **0.003** |
| **accuracy\*block number\*trial number (within a block)** | -0.002 | [-0.024, 0.019] | 0.835 |
| **observation\*block number\*trial number (within a block)** | -0.001 | [-0.023, 0.02] | 0.903 |
| **accuracy\*observation\*block number\*trial number (within a block)** | 0.003 | [-0.019, 0.024] | 0.816 |

*Note. This model tested whether ERP amplitudes (ERN/CRN) varied as a function of accuracy (correct vs. error), observation condition (nonsocial vs. social), block number, and trial number within a block. Full model results, including standardized beta coefficients, confidence intervals, and p-values, are presented. Significant effects in bold font.*

### Trial-level behavioral outcomes as a function of accuracy, observation, block number, and trial number

Having identified a three-way interaction between accuracy, observation condition, and trial number (within a block) predicting trial-level ERN/CRN, as well as a main effect of block on ERN/CRN, we explored whether similar effects were present in the (incongruent) trial-level behavioral data. One model tested the prediction of trial-level accuracy-outcomes from observation condition, block number, trial number within a block, and their interactions. A similar model was fit to test prediction of trial-level RT from accuracy, observation condition, block number, trial number within a block, and their interactions. Consistent with the presence of an overall vigilance decrement across blocks, we observed a significant negative association between block number and accuracy outcomes (β = -0.187, 95% CI [-0.234, -0.140], *p* < 0.001), such that as blocks progressed, accuracy decreased. A significant main effect of observation condition was also present, with higher accuracy outcomes in the nonsocial condition overall (β = -0.082, 95% CI [-0.130, -0.034], *p* < 0.001). For RT, there was a significant interaction between accuracy and block number (β = -0.033, 95% CI [-0.050, -0.015], *p* < 0.001)—in error trials, RT increased across blocks (β = 0.033, 95% CI [0.0002, 0.065], *p* = 0.049), whereas in correct trials, RT decreased (β = -0.033, 95% CI [-0.046, -0.019], *p* < 0.001). Additionally, a significant interaction between observation condition and block number was found (β = 0.026, 95% CI [0.008, 0.043], *p* = 0.004)—in the social condition, RT increased across blocks (β = 0.026, 95% CI [0.001, 0.050], *p* = 0.038), while in the nonsocial condition, RT decreased across blocks (β = -0.026, 95% CI [-0.051, -0.001], *p* = 0.043). However, no identified effect in the trial-level behavioral data mirrored the three-way interaction between accuracy, observation condition, and trial number (within a block) predicting trial-level ERN/CRN. These data suggest the observed trial-level ERN/CRN effect reported above is unlikely to arise as the result of a behavioral confound impacting trial-level ERP measurement. Detailed results are presented in Supplemental Tables S1 and S2.

### Exploratory analyses testing effects of feedback type and trial-level RT

Next, we sought to rule out the possibility that the observed three-way interaction between accuracy, observation condition, and trial number within a block predicting trial-level ERN/CRN could be explained by variation in the type of feedback provided immediately before each block (i.e., whether "Good Job" or "Respond Faster" was presented). Given that feedback was contingent on performance (following recommended best practice; Gehring et al., 2012), with “Respond More Accurately” feedback present for only 13/40 participants in the nonsocial condition and 10/39 participants in the social condition, this feedback type was excluded from analyses involving feedback type (coded as missing for all participants). Similarly, feedback type for the first block was coded as missing because no feedback preceded the first block. We then fit an LMM similar to our primary analysis reported above, when further including feedback type as a predictor (coded as -1 for "Good Job" and 1 for "Respond Faster"); for parsimony, block number was dropped from this model, as it was not significant in the primary analysis. Thus, we fit an LMM predicting trial-level ERN/CRN from accuracy, observation condition, trial number (within a block), feedback type, and their interactions. Crucially, this LMM revealed that the three-way interaction between accuracy, observation condition, and trial number (within a block) remained significant (β = 0.026, 95% CI [0.002, 0.050], *p* = 0.035), even when controlling for feedback type. Moreover, we observed no evidence supporting the notion that feedback type modulates the reported interaction between accuracy, observation condition, and trial number (within a block) predicting trial-level ERN/CRN, as all effects involving feedback type were not significant (all *p* > .320). Detailed results are presented in Supplemental Table S3.

We also sought to rule out the possibility that the observed three-way interaction among accuracy, observation condition, and trial number within a block predicting trial-level ERN/CRN could have resulted from a behavioral confound (RT) impacting ERP measurement. In this model, trial-level RT, accuracy, observation condition, trial number (within a block), and all interaction terms as fixed effects, with ERP amplitude (ERN/CRN) as the dependent variable. Consistent with our primary findings, we again observed a significant three-way interaction among accuracy, observation condition, and trial number (β = 0.033, 95% CI [0.009, 0.057], p = 0.007). Complete results are reported in Supplemental Table S4.

## Discussion

In the current study, we employed mixed-effects modeling of single-trial ERPs to examine whether social observation influences the trajectory of performance monitoring dynamics across both shorter (within blocks) and longer (between blocks) timescales. Analyses revealed that social observation significantly influenced ERN/CRN trajectories at the shorter timescale: for blocks performed under social observation (but not while alone) ERN magnitudes increased across trials and CRN magnitudes decreased. In contrast, at the longer timescale, we observed significant decreases in both ERN and CRN magnitudes across blocks—consistent with a vigilance decrement. Social observation did not significantly moderate ERN/CRN trajectories at the longer timescale, nor did we observe a significant main effect of social observation, on average. Instead, the impact of social observation exhibited a degree of selectivity within the current study, only significantly influencing ERN/CRN trajectories at the shorter timescale. Collectively, these results highlight the importance of considering trial-by-trial (and block-by-block) variation in the ERN/CRN to fully characterize the impacts of social observation on performance monitoring.

To our knowledge, this is the first study to demonstrate that social observation influences performance monitoring trajectories over a relatively short timescale. The results are broadly consistent with prior work reporting that social observation is associated with either an increase in the ERN (Hajcak et al., 2005a; Kim et al., 2005; Niu et al., 2023; Voegler et al., 2018) or increases in the relative difference between the ERN/CRN (Clayson et al., 2024; Schillinger et al., 2016). However, the trial-by-trial modulation of the ERN (and CRN) observed here extends prior findings by demonstrating that the effects of social observation on performance monitoring are dynamic within blocks. Importantly, social observation exhibited a differential influence on within-block ERN and CRN trajectories—the ERN increased in magnitude (became more negative), whereas the CRN decreased in magnitude (became less negative) across trials within a block. Prior work suggests that a larger (more negative) ERN is at least partially reflective of error-specific processing that scales with the motivational significance of errors (Gehring et al., 2012; Hajcak, Moser, Yeung, & Simons, 2005b; Franck Vidal et al., 2021). In contrast, a larger (more negative) CRN may index increased uncertainty and/or misclassifying correct trials as errors (Pailing & Segalowitz, 2004; Scheffers & Coles, 2000). Drawing on these ideas, increases in the ERN can be taken as evidence for enhanced motivational significance of errors. Similarly, increases in the relative difference between ERN/CRN can be taken as evidence for increased fidelity of performance monitoring, given more accurate signaling of error (ERN) and correct (CRN) responses. Taken together, our results suggest that social observation elicits within-block increases in *both* the motivational significance of errors (significant negative ERN slope), as well as increases in the fidelity of performance monitoring (significant differences between the negative ERN slope and positive CRN slope). This interpretation is consistent with the notion that errors made in the presence of others are viewed as more consequential, due to potential social judgment (Van Meel & Van Heijningen, 2010).

Exploratory analyses identified no evidence to suggest that the effect of social observation on within-block ERN/CRN trajectories could be explained by possible confounds of behavioral responding or block-wise feedback. Specifically, exploratory mixed-effects models predicting trial-level ERN/CRN identified the same significant three-way interaction between accuracy, social observation, and trial number (within blocks), even when controlling for trial-level RT or block-level feedback type. Similarly, mixed-effects models predicting trial-level RT and accuracy outcomes revealed no evidence for an effect of social observation on trial-by-trial behavioral trajectories that could account for the reported ERN/CRN effect.

At the longer timescale (across blocks), both the ERN and CRN exhibited a general, non-specific reduction in magnitude across blocks in both the social and nonsocial conditions— a pattern consistent with a vigilance decrement (fatigue effect) and which replicates prior findings (Boksem et al., 2006; Bonnefond et al., 2011; Clayson et al., 2025; Kato et al., 2009; Li, Wang, & Li, 2022; Pattyn, Neyt, Henderickx, & Soetens, 2008; Volpert-Esmond et al., 2018).

However, contrary to our hypothesis, we found no evidence that social observation counteracted such vigilance decrements by sustaining or amplifying ERN magnitudes (or the relative ERN/CRN difference) at the longer timescale (across blocks). Some prior evidence suggests that the effects of vigilance decrements on performance monitoring may be modulated by motivational factors (Boksem et al., 2006; Bonnefond et al., 2011). For example, a mix of monetary incentives and social comparison has been shown to partially buffer against fatigue-related reductions in ERN amplitude over time (Boksem et al., 2006). To our knowledge, however, the present study is the first to test whether social observation (as opposed to social comparisons or monetary incentives) can similarly modulate the effects of fatigue on ERN trajectories at a relatively long timescale (across blocks). Preliminary evidence from Bonnefond et al. (2011), which analyzed data from 18 participants in a between-subjects design (9 per group), suggested that social observation may modulate longer-term CRN trajectories.

However, as their study did not examine ERN trajectories over time, it is difficult to directly compare their findings with results of the current study. CRN decreases that co-occur with ERN decreases indicate broad disengagement of task-related performance monitoring (e.g., due to fatigue), consistent with the pattern that we observed at the longer timescale (across blocks). Conversely, CRN decreases that co-occur with a stable/increasing ERN (i.e., an increase in the relative ERN/CRN difference) indicate enhancements in task-related performance monitoring, consistent with the effect of social observation that we observed at the shorter timescale (within blocks). Ultimately, our data replicate prior evidence for a broad vigilance decrement present for both the ERN and CRN across blocks (Boksem et al., 2006; Kato et al., 2009; Lorist et al., 2005; Volpert-Esmond et al., 2018; Xiao et al., 2015), and we do not find evidence that social observation impacts ERN or CRN trajectories across blocks (only across trials within blocks). More work is needed to further test possible effects of social observation on the ERN/CRN over longer timescales, including as a function of individual differences in motivation or affect.

In the current study, the effect of social context on ERN/CRN trajectories was specific to the shorter timescale (within blocks), and we did not detect a significant main effect of social observation overall, nor an effect of social observation on ERN/CRN trajectories at the longer timescale (across blocks). The lack of a significant main effect of social observation is inconsistent with several prior studies reporting that the trial-averaged ERN (or the relative ERN/CRN difference) is larger under social observation (Clayson et al., 2024; Hajcak et al., 2005a; Kim et al., 2005; Niu et al., 2023; Schillinger et al., 2016; Voegler et al., 2018). One possibility is that the lack of a significant main effect of social observation, as well as the absence of social observation effects at the longer timescale, relates to the number of trials per block in our task (40 trials/block). Within blocks, we found that as trials progressed, social observation was associated with a pattern of increasing ERN amplitudes, as well as increases in the relative difference between ERN/CRN. If similar ERN/CRN trajectories are present for longer blocks, then trial averages based on longer blocks would contain a greater proportion of relatively larger ERN/CRN magnitudes; this might increase the likelihood of detecting either an overall main effect of social observation, or an effect of social observation at longer timescales (across blocks). Re-analyses of prior social observation studies that employ alternative block lengths could be particularly informative.

Another possible reason that we observed no main effect of social observation on ERN/CRN, and no effect of social observation at the longer timescale, could relate to the method of social observation employed in our study. Participants were observed remotely by a second experimenter via Zoom, with a stand-mounted iPad positioned outside the participant’s direct line of sight. It is possible that the limited visibility and physical presence of the iPad observer reduced the salience or perceived evaluative impact of the social manipulation. Compared to studies that used in-person observers, our setup may have represented a more subtle form of social evaluation and evoked less intense social pressure. Nevertheless, the current study revealed that social observation was associated with dynamic changes in ERN and CRN across trials within each block. This pattern suggests that the influence of social evaluation on performance monitoring may not operate as a uniform, condition-wide shift in neural responses, but rather unfolds dynamically over time. Trial-averaged ERP approaches, which aggregate across all trials within a condition, would likely obscure these dynamic patterns (Heise, Mon, & Bowman, 2022b; Volpert-Esmond et al., 2018). In contrast, trial-level analyses allow for the detection of temporally specific effects—such as the gradual divergence of ERN and CRN amplitudes across trials within a block—that provides a richer understanding of how social observation impacts performance monitoring across time.

### Limitations and future directions

The current study is not without limitations. We had observers read the block feedback aloud within the social condition, to remind participants of the social observation. Thus, it is possible that presenting feedback vocally vs. silently could have impacted participants’ responsivity to feedback, as opposed to resulting from social observation alone. Exploratory analyses testing whether feedback type and social observation interacted to predict ERN/CRN were not significant and are thus inconsistent with this interpretation. Nonetheless, future work that directly manipulates the method of feedback delivery, as well as feedback content, would be informative. Prior work demonstrates that the effect of social observation on ERN/CRN is influenced by individual differences (Buzzell et al., 2017; Niu et al., 2023). Having established here that social observation influences within-block ERN/CRN trajectories, future work could leverage a larger sample size to investigate whether individual differences—such as trait anxiety, social sensitivity, or cognitive control—moderate the trajectory of neural responses under social observation.

## Conclusions

The present study used a within-subject design and mixed-effects modeling of single-trial ERPs to investigate whether social observation influences the trajectory of performance monitoring dynamics across both shorter and longer timescales. We found that, in the social condition (but not the nonsocial condition) the pattern of ERN and CRN changed over the shorter timescale (across trials within a block). Specifically, social observation was associated with ERN magnitudes becoming increasingly larger (more negative), while CRN became increasingly smaller (less negative) as trials progressed within a block. We did not observe an overall increase in ERN amplitude in the social (vs. nonsocial) condition, nor did we find evidence that social observation influences ERN/CRN trajectories at the longer timescale (across blocks). Instead, at the longer timescale, both the ERN and CRN declined in magnitude over time in both conditions—a pattern consistent with a general vigilance decrement (fatigue effect) over the course of a prolonged task. To our knowledge, this is the first study to examine how social observation modulates performance monitoring trajectories at both shorter (within blocks) and longer (across blocks) timescales, highlighting the value of multilevel modeling approaches for uncovering dynamic fluctuations in performance monitoring that may otherwise be obscured by condition-averaged analyses. Moreover, these data highlight the importance of considering task structure and timescale when investigating how motivational contexts like social observation modulate performance monitoring processes. The current findings lay important groundwork for future research investigating how individual differences—such as those related to motivational states, affective traits, or cognitive control—may shape the temporal dynamics of performance monitoring.

## Acknowledgements

Funding Research reported in this publication was supported by the National Institute of Mental Health of the National Institutes of Health under award number R01MH131637.

## Availability of data and materials

Deidentified data are available from the corresponding author upon request.

## Code availability

The Psychopy task and data processing scripts are publicly available on the following GitHub repository: https://github.com/NDCLab/social-flanker-error.

