## Supplementary for "Social observation influences the trajectory of performance monitoring across trials: evidence from single-trial estimates of the ERN and CRN"

Supplementary Materials for
Social evaluation elicits a trajectory of increasing error monitoring across trials: evidence from trial-level estimates of the ERN

### EEG Preprocessing Details

EEG data were preprocessed using MATLAB R2021b (MathWorks Inc., Sherborn, MA, USA), the EEGLAB toolbox, and a modified version of the MADE pipeline (Debnath et al., 2020; Delorme & Makeig, 2004). Specifically, data were filtered via a high-pass filter at 0.1 Hz, followed by a low-pass filter with a passband of 49 Hz and a stopband of 59 Hz. The FASTER EEGLAB plugin (Nolan, Whelan, & Reilly, 2010) was used to identify/exclude bad channels. To further remove noise components originating from ocular and/or muscular activity, we performed independent component analysis (ICA). To improve ICA decomposition, a copy of each dataset was made, high-pass filtered at 1 Hz, and segmented into 1-second epochs (Debnath et al., 2020). If a channel had more than 20% artifacted epochs, that channel was removed from both the copied and original dataset (Debnath et al., 2020). No more than 10% of channels were identified as bad for all participants. ICA was then performed on the filtered/cleaned dataset. ICA weights were copied back to the original dataset and the adjusted-ADJUST algorithm (Leach et al., 2020; Mognon, Jovicich, Bruzzone, & Buiatti, 2011) used to identify independent components (ICs) corresponding to artifacts; identified ICs were subtracted from the data. Data were segmented into 3-second epochs (-1 to 2 seconds, relative to the response) and the DC-offset subtracted from each epoch. To eliminate any remaining ocular artifacts, epochs were rejected if a ±125 μV threshold was exceeded for channels located near the eyes. For all other channels not located near the eyes, if the ±125 μV threshold was exceeded, the channel was interpolated (unless more than 10% of channels in a given epoch exceeded the threshold, in which case the entire epoch was removed). A spherical spline interpolation was utilized to interpolate missing channels, after which channels were re-referenced to the average of all channels. Finally, the (-400 to -200) pre-response baseline was subtracted from each epoch.

Table S1. Linear mixed-effects model of trial-level accuracy as a function of observation condition, block number, trial number within a block, and their interactions.

| Variables | beta | beta [95% CI] | *p* |
|---|---|---|---|
| **observation** | ***-0.082** | **[-0.130, -0.034]** | **< 0.001** |
| **block number** | ***-0.187** | **[-0.234, -0.140]** | **< 0.001** |
| **trial number (within a block)** | -0.023 | [-0.070, 0.025] | 0.347 |
| **observation*block number** | 0.035 | [-0.012, 0.082] | 0.140 |
| **observation*trial number (within a block)** | -0.004 | [-0.051, 0.043] | 0.877 |
| **block number*trial number (within a block)** | -0.038 | [-0.085, 0.009] | 0.109 |
| **observation*block number*trial number (within a block)** | -0.040 | [-0.087, 0.007] | 0.097 |

Table S2. Linear mixed-effects model of trial-level reaction time as a function of accuracy, observation condition, block number, trial number within a block, and their interactions.

| **Variables** | **beta** | **beta [95% CI]** | ***p*** |
|---|---|---|---|
| **accuracy** | ***0.441** | **[0.423, 0.459]** | **< 0.001** |
| **observation** | 0.002 | [-0.016, 0.019] | 0.858 |
| **block number** | -0.000 | [-0.018, 0.017] | 0.994 |
| **trial number (within a block)** | -0.005 | [-0.022, 0.012] | 0.576 |
| **accuracy*observation** | -0.011 | [-0.029, 0.006] | 0.196 |
| **accuracy*block number** | ***-0.033** | **[-0.050, -0.015]** | **< 0.001** |
| **observation*block number** | ***0.026** | **[0.008, 0.043]** | **0.004** |
| **accuracy*trial number (within a block)** | -0.007 | [-0.024, 0.011] | 0.447 |
| **observation*trial number (within a block)** | 0.014 | [-0.003, 0.031] | 0.108 |
| **block number*trial number (within a block)** | 0.009 | [-0.008, 0.027] | 0.297 |
| **accuracy*observation*block number** | -0.007 | [-0.024, 0.010] | 0.431 |
| **accuracy*observation*trial number (within a block)** | -0.011 | [-0.028, 0.006] | 0.217 |
| **accuracy*block number*trial number (within a block)** | 0.002 | [-0.015, 0.020] | 0.814 |
| **observation*block number*trial number (within a block)** | 0.012 | [-0.005, 0.030] | 0.170 |
| **accuracy*observation*block number*trial number (within a block)** | 0.006 | [-0.024, 0.011] | 0.498 |

Table S3. Linear mixed-effects model examining whether feedback type modulates the interaction between accuracy, observation condition, and trial number on ERN/CRN amplitudes. Feedback type was coded as “Good Job” = -1 and “Respond Faster” = 1. In this model, feedback type, accuracy, observation condition, trial number (within a block), and all interaction terms as fixed effects, with ERP amplitude (ERN/CRN) as the dependent variable.

| **Variables** | **beta** | **beta [95% CI]** | ***p*** |
|---|---|---|---|
| **accuracy** | ***0.262** | **[0.238, 0.287]** | **< 0.001** |
| **observation** | -0.007 | [-0.032, 0.017] | 0.553 |
| **trial number (within a block)** | -0.004 | [-0.028, 0.020] | 0.745 |
| **go_faster** | -0.011 | [-0.038, 0.016] | 0.414 |
| **accuracy*observation** | -0.008 | [-0.032, 0.016] | 0.516 |
| **accuracy*trial number (within a block)** | 0.023 | [-0.001, 0.048] | 0.061 |
| **observation*trial number (within a block)** | -0.002 | [-0.026, 0.023] | 0.896 |
| **accuracy*go_faster** | 0.001 | [-0.023, 0.026] | 0.914 |
| **observation*go_faster** | 0.012 | [-0.012, 0.037] | 0.321 |
| **trial number (within a block)*go_faster** | 0.001 | [-0.024, 0.025] | 0.954 |
| **accuracy*observation*trial number (within a block)** | ***0.026** | **[0.002, 0.050]** | **0.035** |
| **accuracy*observation*go_faster** | -0.004 | [-0.028, 0.020] | 0.732 |
| **accuracy*trial number (within a block)*go_faster** | 0.010 | [-0.014, 0.035] | 0.400 |
| **observation*trial number (within a block)*go_faster** | -0.004 | [-0.029, 0.020] | 0.720 |
| **accuracy*observation*trial number (within a block)*go_faster** | 0.006 | [-0.018, 0.030] | 0.635 |

Table S4. Linear mixed-effects model examining whether reaction time modulates the interaction between accuracy, observation condition, and trial number on ERN/CRN amplitudes. In this model, trial-level RT, accuracy, observation condition, trial number (within a block), and all interaction terms as fixed effects, with ERP amplitude (ERN/CRN) as the dependent variable.

| **Variables** | **beta** | **beta [95% CI]** | ***p*** |
|---|---|---|---|
| **accuracy** | ***0.222** | **[0.196, 0.248]** | **< 0.001** |
| **observation** | -0.005 | [-0.03, 0.02] | 0.678 |
| **trial number (within a block)** | -0.002 | [-0.026, 0.022] | 0.858 |
| **log_rt** | 0.013 | [-0.009, 0.035] | 0.255 |
| **accuracy*observation** | -0.008 | [-0.033, 0.017] | 0.525 |
| **accuracy*trial number (within a block)** | 0.014 | [-0.01, 0.038] | 0.258 |
| **observation*trial number (within a block)** | -0.013 | [-0.037, 0.01] | 0.273 |
| **accuracy*log_rt** | ***-0.081** | **[-0.1, -0.061]** | **< 0.001** |
| **observation*log_rt** | 0.006 | [-0.014, 0.025] | 0.569 |
| **trial number (within a block)*log_rt** | 0.016 | [-0.003, 0.034] | 0.096 |
| **accuracy*observation*trial number (within a block)** | ***0.033** | **[0.009, 0.057]** | **0.007** |
| **accuracy*observation*log_rt** | -0.008 | [-0.027, 0.011] | 0.396 |
| **accuracy*trial number (within a block)*log_rt** | 0.015 | [-0.003, 0.033] | 0.113 |
| **observation*trial number (within a block)*log_rt** | 0.005 | [-0.014, 0.023] | 0.619 |
| **accuracy*observation*trial number (within a block)*log_rt** | 0.005 | [-0.013, 0.024] | 0.567 |
